# A resource for the comparison and integration of heterogeneous microbiome networks

**DOI:** 10.1101/2022.08.07.503059

**Authors:** Zhenjun Hu, Dileep Kishore, Yan Wang, Gabriel Birzu, Charles DeLisi, Kirill Korolev, Daniel Segrè

## Abstract

Naturally occurring microbial communities often comprise thousands of taxa involved in complex networks of interactions. These interactions can be mediated by several mechanisms, including the competition for resources, the exchange of signals and nutrients, cell-cell contact and antibiotic warfare. In addition to direct measurements and computational predictions of interactions, abundant data on microbial co-occurrence associations can be inferred from correlations of taxa across samples, which can be estimated from metagenomic, and amplicon datasets. The analysis and interpretation of interaction and correlation networks are limited by the challenge of comparing across different datasets, due to heterogeneity of the data itself and to the lack of a platform to facilitate such comparisons. Here, we introduce the Microbial Interaction Network Database (MIND) - a web-based platform for the integrative analysis of different types of microbial networks, freely available at http://microbialnet.org/. In addition to containing a growing body of curated data, including amplicon-based co-occurrence networks, genome-scale model-derived networks, metabolic influence networks and horizontal gene transfer networks, MIND allows users to upload and analyze newly generated networks using a JSON format and standard NCBI taxonomy. The platform provides convenient functions to compare and query multiple networks simultaneously, and to visualize and export networks and datasets. Through some illustrative examples, we demonstrate how the platform might facilitate discoveries and help generate new hypotheses on host-associated and environmentally important microbial ecosystems through the power of knowledge integration.

## Introduction

Microbial communities play a major role in global biogeochemical cycles [1–5], human health [6–9], and future environmental sustainability [10–13]. Understanding these communities and being able to steer them towards desired states are therefore important goals at the crossroads of microbiology, synthetic biology, microbial ecology and metabolic engineering. These tasks are arduous, as natural microbial ecosystems often involve thousands of distinct microbial taxa, whose destinies are tightly coupled with each other [14–16]. The interdependencies between microbes in communities, and the intracellular wirings that drive these ecological webs are key determinants of community structure and dynamics [17–19]. Thus, being able to catalog, analyze and compare inter-microbial networks across datasets and different types of interactions could have a great impact on our capacity to understand and control communities.

The overwhelming majority of microbe-microbe networks published in conjunction with microbiome studies are correlation networks, in which an edge between two taxa encodes the fact that the presence of one taxon in a sample is indicative of an increased chance of the presence or absence of another. Correlations are generally inferred from 16S rRNA amplicon or metagenomic sequencing data, using an expanding and diverse set of specialized computational tools [20–23]. Despite widespread usage of correlation networks, interpretability is limited due to several factors, including the compositional and noisy nature of microbiome data generally used to infer co-occurrence networks, the lack of a single agreed upon approach for correlation estimation [23], and the inherent poor understanding of how correlations relate to actual interactions. Additionally, comparing across different publicly available correlation networks is currently extremely challenging, due to the fact that datasets often sit on distinct servers, use different formats or are only available as supplementary tables in separate individual papers [24].

In addition to correlation networks, an increasing number of experimental and computational efforts are specifically focused on assessing actual microbe-microbe interdependencies or direct interactions, e.g., by determining how the growth of an organism, or the metabolites it uses and secretes are modified by the presence of another organism. These types of data are extremely diverse and span many different types of experimental setup and computational approaches. They include Lotka-Volterra coefficients describing inter-species influences, inferred from longitudinal data [25–28]; direct measurements of how pairs of species affect each other’s growth in well mixed [29–31] or spatially structured environments [19,32–34]; microfluidic droplet co-culture measurements [30,35,36]; monitoring of temporal correlations through single-cell counting in microcosm experiments [37,38]; estimates of degree of horizontal gene transfer across species based on genomic analyses [39,40]; microscopy observation of spatial co-localization of different bacterial species [41–43]; genome-scale model predictions of metabolic competition and cross-feeding [14,44–48]; putative interactions inferred through machine learning approaches from existing datasets [49]; and - last but not least - a myriad of individually reported detailed descriptions of interactions published in peer-reviewed papers [24].

The availability of multiple interaction and correlation networks on a unified platform would enable cross-data comparisons and would facilitate the mining of associations and subnetworks, creating multiple new opportunities for discovery in microbiome research. Similar to prior analyses of genetic networks or protein-interaction networks, integration of microbial interaction networks could spark new discovery and increased understanding of microbiome structure and function. Although a few tools are becoming available for the visual analysis of microbial networks [50–52], to our knowledge no tools exist that can represent, compare and integrate diverse sets of microbial interaction types.

Here we present the Microbial Interaction Network Database (MIND), a prototype for a multi-level integrative software platform for the analysis of large, heterogeneous datasets on microbial inter-dependencies. Note that the current database includes both correlation networks and different types of inter-dependencies and interactions, such as horizontal gene transfer networks, genome-scale model-derived networks, and ecological networks compiled from experimental data.

Throughout this paper, and in the database itself, we will refer broadly to all microbe-microbe networks as “interaction networks”. Our platform combines knowledge from large-scale data-driven statistical inference with mechanistic understanding of how microbes interact with each other. MIND, which is freely accessible at http://microbialnet.org/, is implemented with a service-oriented architecture (SOA) [53] that is an enhanced version of VisANT-Predictome’s three-tier system [54–56], composed of a database, an application server and a web client.

Through MIND, we hope to make it possible for a large number of users to take advantage of multiple datasets existing in the database, for integrative studies of microbial ecosystem structure. We expect that MIND, and its possible extensions, will help capture in a flexible way the complex nature of microbial interdependencies.

In this paper, we describe in detail the functionalities of the MIND platform and provide examples of its potential as a tool for alleviating the data-to-knowledge bottleneck. To exemplify the value of MIND for microbiome research and discovery, we provide a few examples of how one can use the resource to find common interactions among datasets, and compare patterns of connectivity, abundance and co-occurrence. We conclude with a case study focused on the re-analysis of colorectal cancer data, where, in addition to identifying robust network modules, we show how one can use MIND to explore possible connections with the oral microbiome. While MIND currently contains mostly human microbiome datasets, it is designed to serve as a broader analysis tool, which can help map microbial interdependencies within and across different biomes.

## Results

### The MIND Database and Web Platform

MIND stores microbial interaction networks and related data from diverse sources. The structure of the database is an extension of the Predictome database [55,57], which permits easy integration of interactions at different scales - proteins, pathways, cells and organisms [56]. Microbes are uniquely identified by their NCBI taxonomy ID [58,59]. Interactions or correlations reported for a collection of microbes under a given set of conditions form a network, and networks are organized using specific contexts that stores abundant metadata, including information about the environment, disease relevance (if appropriate), and experimental and computational methods used to infer interactions. Many of the networks currently present in the database are either curated manually from published studies or generated by a recently developed standardized pipeline (MiCoNE, [23]) applied to published 16S rRNA sequence data. New networks may also be uploaded by users. The abundance profile of different taxa, if available for a specific dataset, can also be stored in the database, and visualized on top of the network structures. Detailed information about database design and implementation can be found in Materials and Methods.

To power the visual analysis of these networks, MIND-web is built as a SPA (Single Page Application) [60] using cutting-edge web technology (d3 [61,62], Bootstrap [63] and AngularJS [64,65]), with standard user conventions and a responsive user interface to support mobile platforms. The platform provides rich functions of interaction queries and exploratory navigation for the microbes of interest, and comparative analysis of the networks from different sources (Fig. S1). In addition, it provides functions that are specific to microbial networks, such as level-up approximation to transform the network nodes to a specified higher-level taxonomic rank (see Materials and Methods).

### Exploration of microbial interactions

#### Interaction visualization

A microbial interaction is generally represented as an edge (line) connecting two nodes (microbes; Fig. 1A). The edge thickness is proportional to the weight of the interaction, which may have different meanings for different types of networks. A pair of microbes will be connected by multiple edges when the interaction appears in different contexts (see next section for more information about context); these are distinguished by different edge colors (Fig.1B,C). Positive and negative associations are represented by different line types, with negative associations in particular, represented by dashed lines. By default, the node color maps to a microbe’s taxonomy ranking.

**Figure 1.**
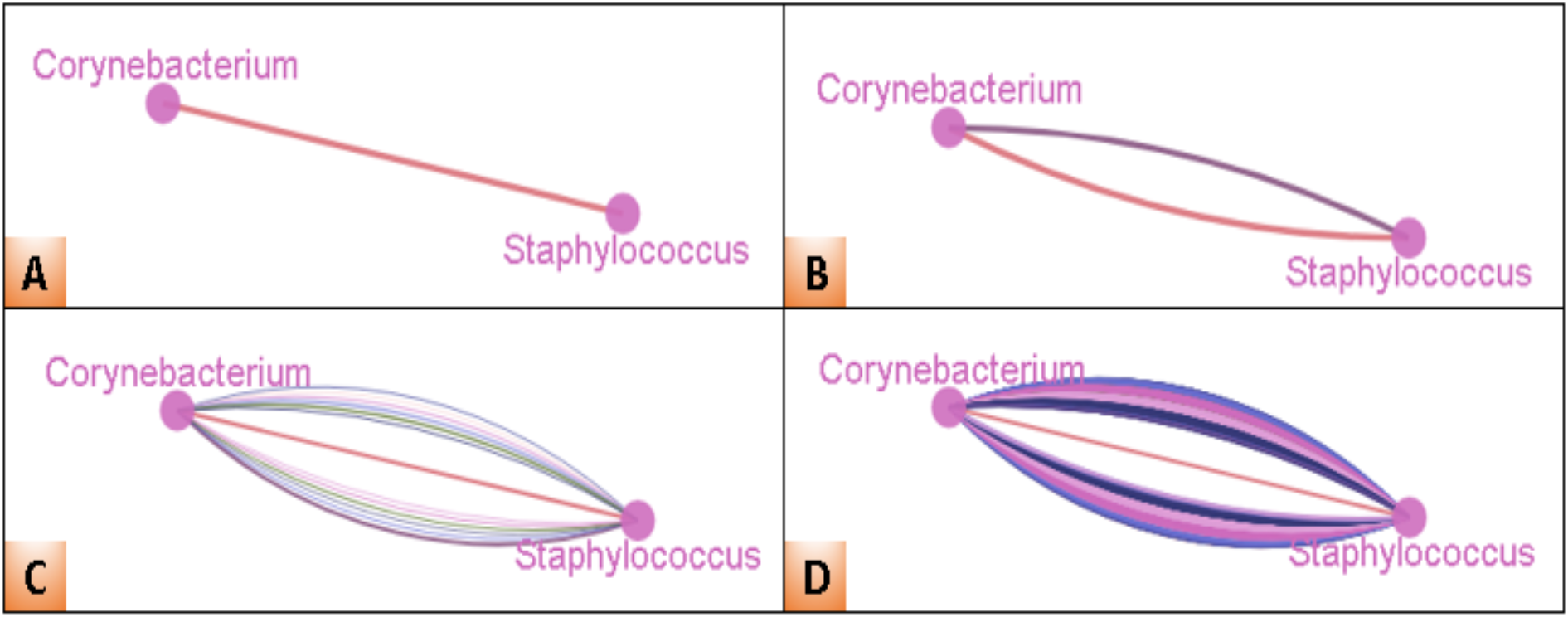
Visualization of microbial interactions. A) Corynebacterium and Staphylococcus are correlated in the Crohn’s disease samples from the Swiss IBD cohort [67]. B) This interaction can be reproduced in the Crohn’s disease samples from the Bern gastroenterology clinics cohort [67]. The two contexts are distinguished by edge color. C) The same pair of microbes are connected by 13 edges (13 contexts) reported by MIND. D) There are a total of 819 interactions between Corynebacterium and Staphylococcus, either in different contexts or the same context with different weights reported by MIND.

MIND not only enables easy mining and display of microbial interactions in a given context (Fig. 1A) but also facilitates visualization of interactions for all known contexts (Fig. 1C). The same pair of nodes can be linked by multiple edges. For example, since microbes at a given taxonomic level are generally represented by Operational Taxonomic Units (OTUs) in 16S data, it is not unusual to find multiple correlations with different weights between a pair of nodes in the same context (multiple OTUs may be mapped to same species/genus). In such a case, a user has the option of combining duplicated interactions, as shown in Fig. S2. By default, these are merged, with the weights of the final edges computed as averages of the duplicate ones (Fig.1C). As an example, Fig. 1D displays a total of 819 correlations in 13 contexts for the pair of microbes shown in Fig. 1C.

Detailed information about a node or edge can be obtained using MIND’s mouse-over feature. Zooming and panning can be achieved using the mouse wheel and dragging. Step-wise zooming is also provided through buttons on the toolbox panel. In addition, the MIND platform provides rich customization functions for both nodes and edges, including color, size and transparency. The platform also implements an optimized arrangement of node labels to reduce the overlap between labels and nodes/edges, as shown in Fig. S3.

#### Exploratory navigation

One unique feature of the MIND platform is its support of exploratory navigation to walk through interactions based on one or a few initial microbes. Users can search for microbes of interest by typing a few letters, and the search box will provide a list of matched microbes. The search box also supports copy/paste for comma-delimited lists of microbial names or taxonomy IDs. Once the query finishes, information about the nodes and edges can be obtained by mouse-over. Double-clicking a node will initiate a query of the interactions of the corresponding microbe, which allow the user to walk through the interactions step-by-step. An example of exploratory navigation is shown in Fig. 2.

**Figure 2.**
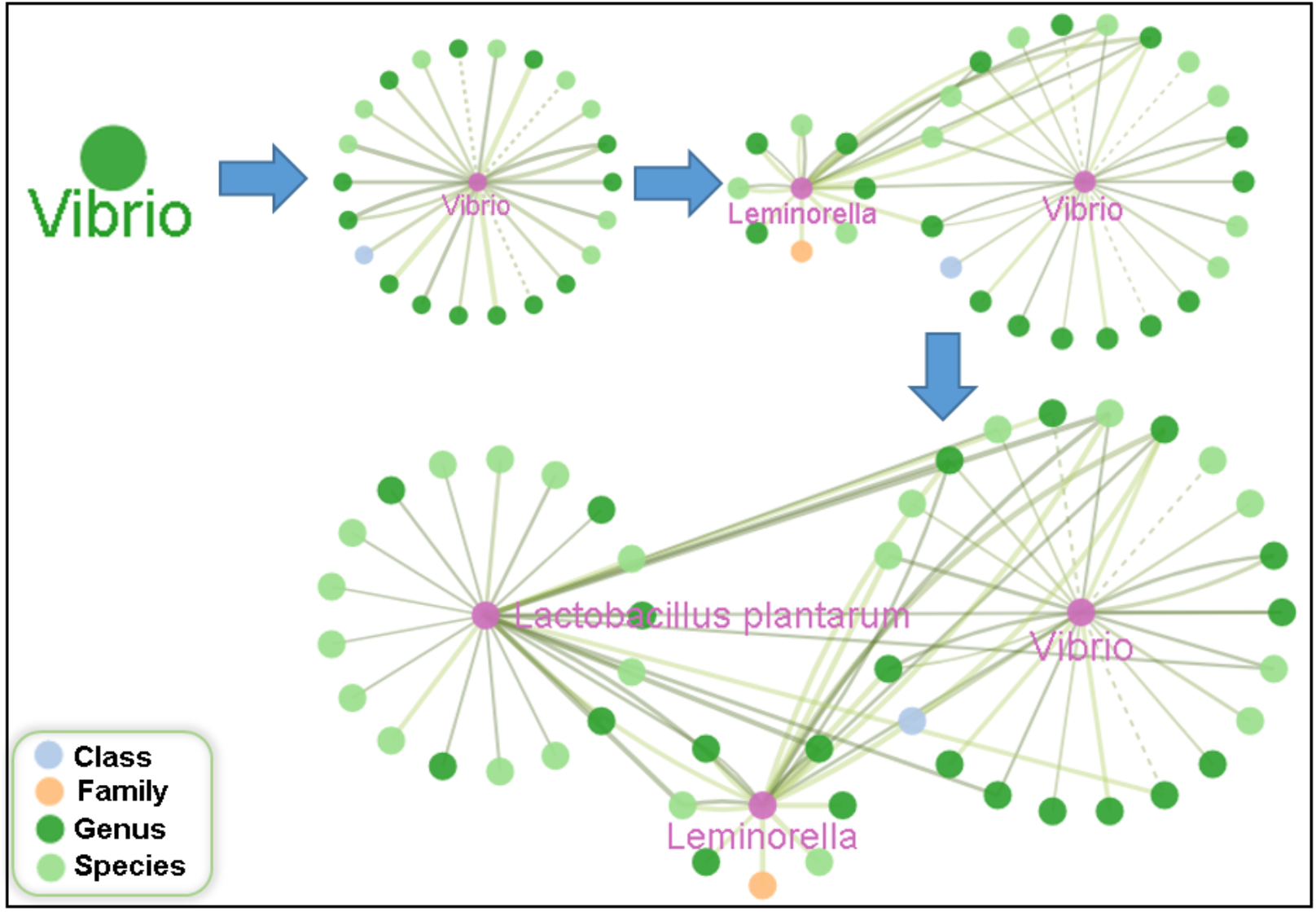
Exploratory navigation of microbial interactions allows quick expansion of the network. The navigation starts by double-clicking the node *Vibrio* followed by *Leminorella* and *Lactobacillus plantarum*. The solid line represents positive correlation while the dashed line represents a negative one. The nodes are colored purple when their interactions have been queried against the database.

#### Network filtering

MIND’s microbial networks are often heterogeneous in taxonomic ranks. For example, the networks shown in Fig. 2 involve four ranks, ranging from species to class. It’s easy to filter such networks using the checkbox taxonomy filters in the toolbox panel (Fig. S1) to turn on/off nodes with specific rankings. The platform also supports the filtering of nodes based on information on their habitats. For example, oral microbiome data can carry metadata obtained from the human oral microbial database (HOMD) [66]. Edges may be filtered out by their weights, either globally, or specific to a given context; second, they can be turned on/off according to context; and finally, they may be filtered out based on the number of connections between a pair of nodes. Figure 3 shows an example where we discovered strong correlations that are consistent in four different groups of samples with inflammatory bowel disease (IBD) using these functions.

**Figure 3.**
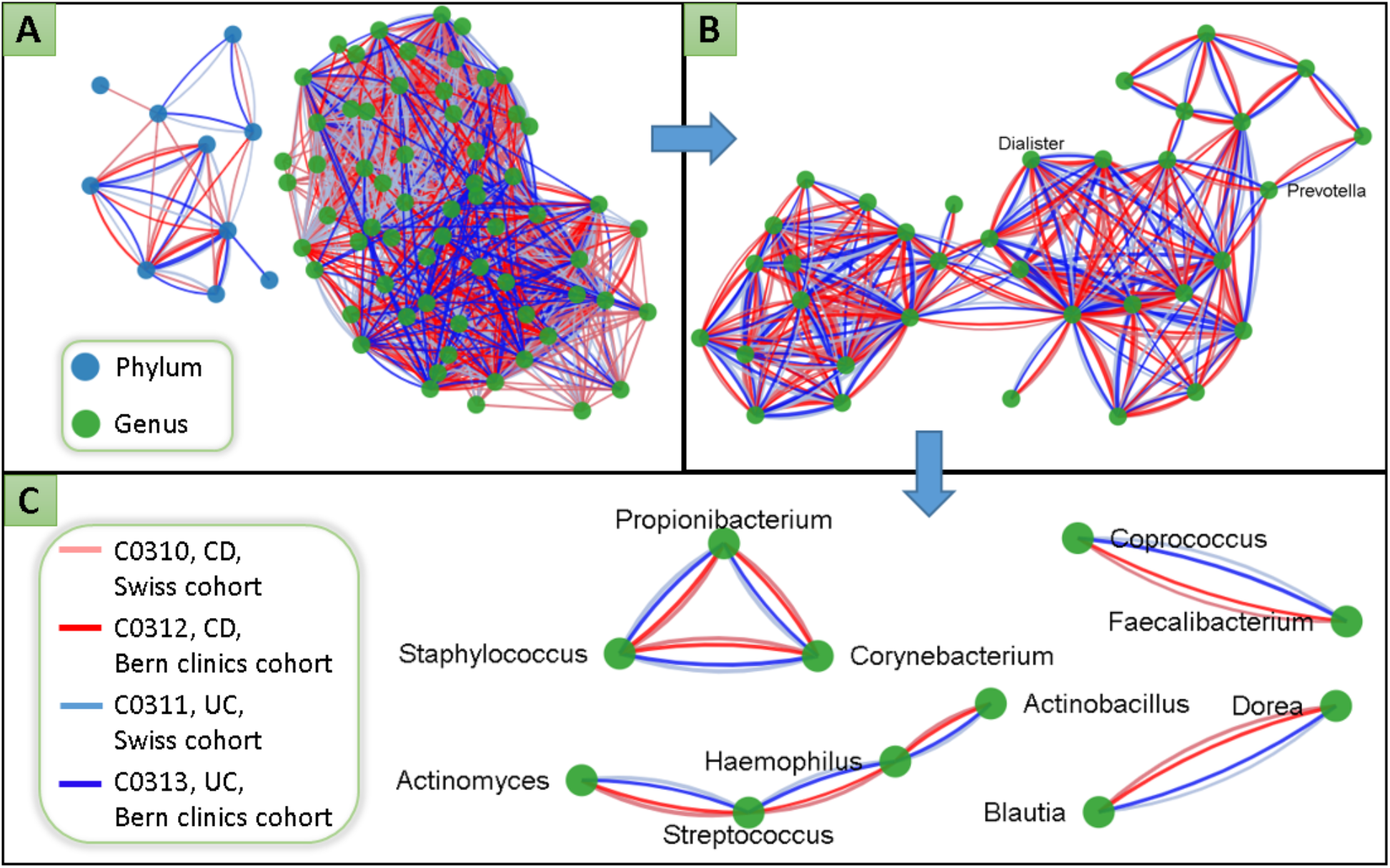
Through a few steps in MIND it is possible to extract, from multiple complex interaction network datasets, a few patterns encoding strong correlations that are consistent across datasets, in this case inflammatory bowel disease (IBD) samples from the Swiss cohort and the Bern clinics cohort [67]. IBD is often phenotypically divided into two types: Crohn’s disease (CD) and ulcerative colitis (UC). Contexts C0310-C0313 are based on the study involving two independent sample cohorts: Swiss IBD cohort and Bern gastroenterology clinics cohort. The MIND analysis steps here consist of: A) Load the four networks of C0310-C0313 by clicking the All button for each context (Fig. S4) and apply force layout (under Layout menu) which will generate a network with two separate modules. B) Continue to filter out nodes with phylum rank, and only keep the links that are consistent in all four contexts by setting 4 as the cutoff for the number of links in the toolbox panel (Fig. S1). Surprisingly the remaining links form one big module. C) Because all four contexts use the Pearson correlation, we apply a global weight cutoff 0.5 to obtain four graphs with strong correlations.

### Navigation of Microbial Interaction Networks (Contexts)

Any microbial network uploaded in MIND can include a set of contexts, i.e., metadata relative to the conditions under which the network was generated, including information about the environment, potential disease relevance, and experimental and computational methods used to infer interactions (Fig. S5). MIND Web provides a set of comprehensive functions to assist users in navigating contexts. As shown in Fig. S1 and S4, each row in the toolbox panel labels a context, and buttons allow users to manipulate the network (detailed Fig. S4), including loading all interactions in a specified context (Fig. 3A), turning the abundance profile on or off, and filtering the interactions with context-specific weight cutoffs. Contexts can be searched using any keywords, and the auto-completion feature makes searching convenient. Three tag lists are provided to filter contexts, including different disease states, interaction types and habitats (Fig. S4).

### Knowledge Integration

In addition to integrated taxonomy knowledge, MIND Web also supports integrated visualization of taxonomic abundances (if available), as well as topological features such as degree distribution. Fig. 4 shows an example where the size of a node is proportional to its degree. Note that nodes in Fig. 4A-D are scaled independently. *Prevotella* which has the highest degree in the integrated network (Fig. 4D), is also most widely distributed across body sites. We have not yet ascertained the extent to which such a correlation can be generalized.

**Figure 4.**
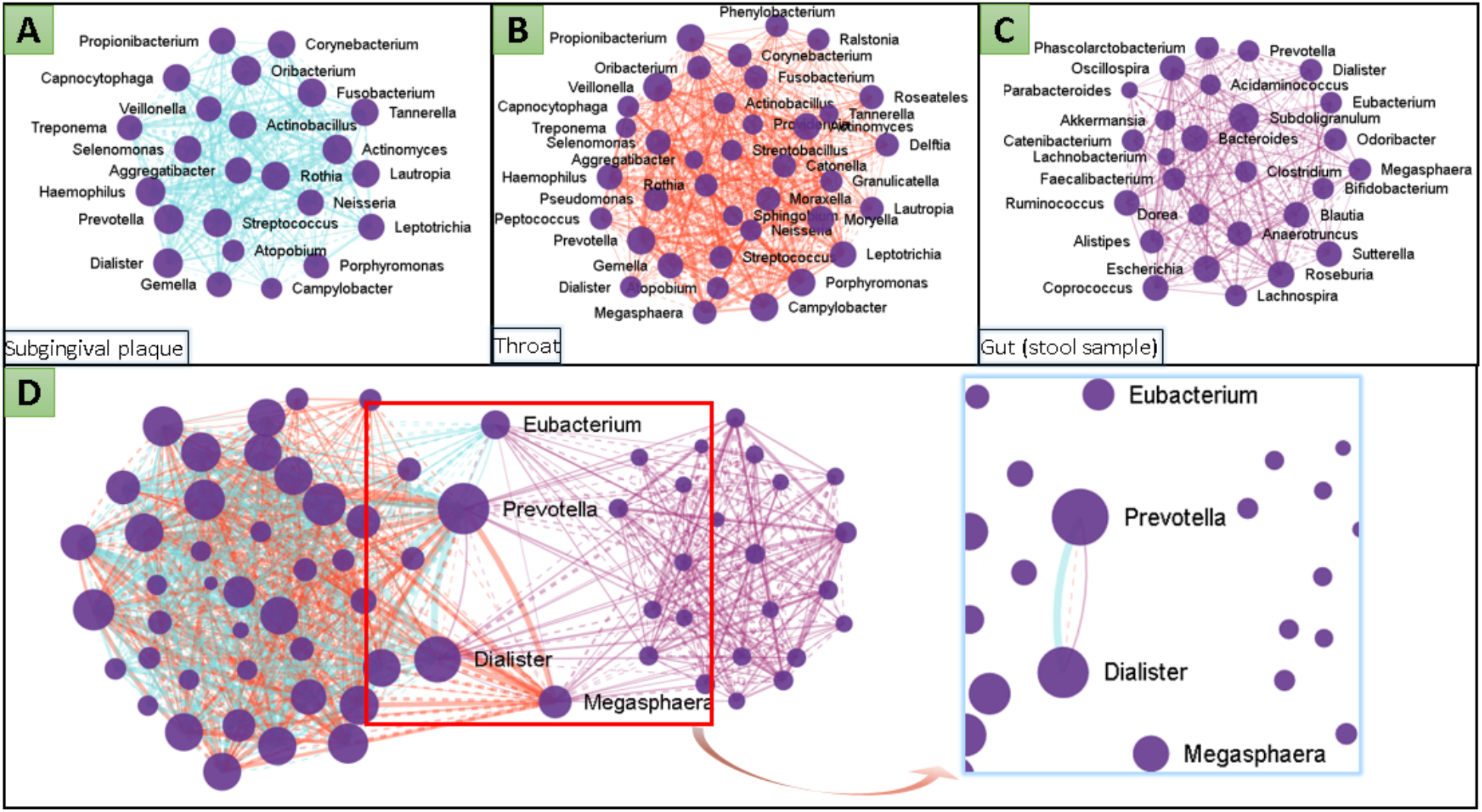
Comparative analysis of microbial co-occurrence networks across different body sites (with edge colors meant to distinguish between the different datasets). The networks are generated using the MIND pipeline based on 16S rRNA sequence data of healthy samples provided by the HMP project [86]. All nodes represent genus, and size is proportional to degree. A) Microbial network at subgingival plaque; B) Microbial network at throat; C) Microbial network at gut (Stool sample); D) integrated microbial network of three body sites, where *Eubacterium, Prevotella, Dialister* and *Megasphaera* serve as keep points to connect oral and gut microbiomes. The zoomed region with the filter set to “3 links” shows that only the interaction between *Prevotella* and *Dialister* is present in all body sites.

Fig. 5 shows examples of abundance visualization over the integrated network where the node size is mapped to its relative abundance, specifically to verify the reproducibility of the abundance between related datasets on IBD (Inflammatory Bowel Disease) microbiome (a Swiss IBD cohort and a Bern gastroenterology clinic cohort [67]. It can be seen that the average microbial abundances for (Crohn’s Disease) CD are similar for the two cohorts (with a slight difference for *Proteobacteria)*, while for ulcerative colitis (UC) there is a significant difference in the abundance for both *Proteobacteria* and *Firmicutes*. On the other hand, the networks of UC are in a better agreement between two cohorts than those of CD.

**Figure 5.**
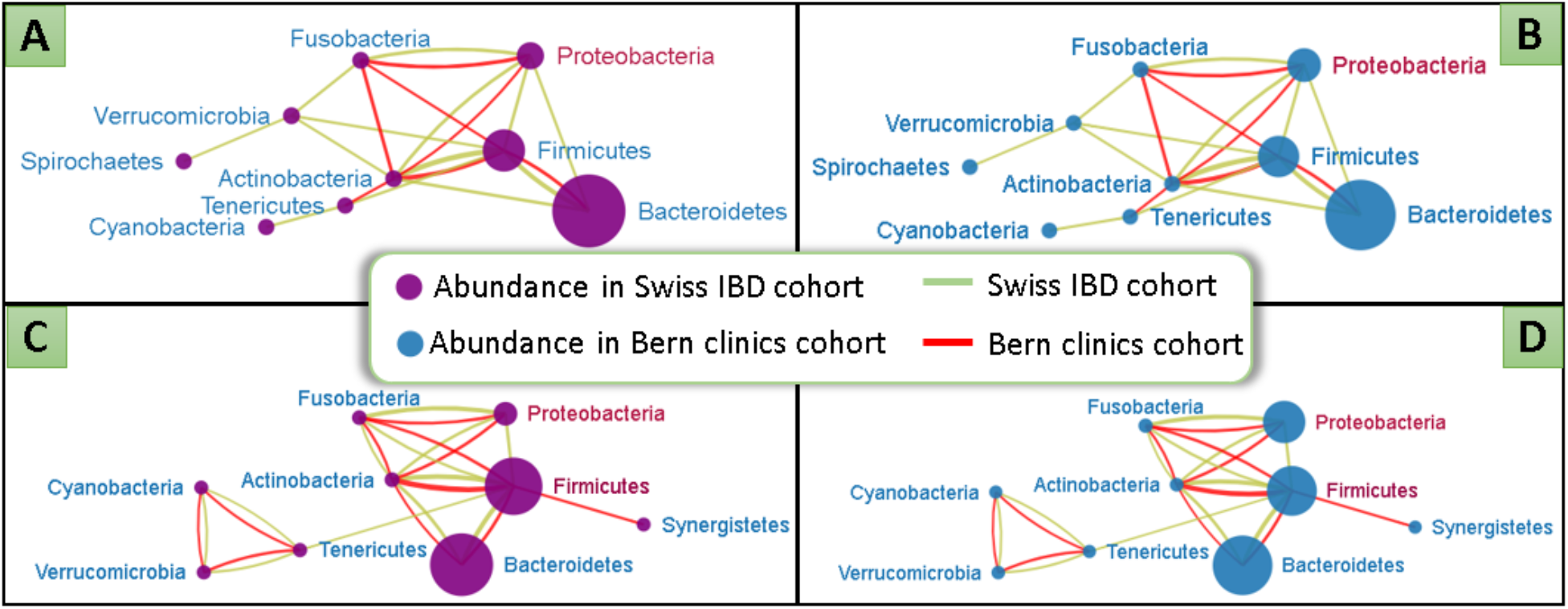
Comparison of abundance profiles between two cohorts for IBD [67], at the level of Phylum. A) The abundance of the Swiss IBD cohort overlaid with an integrated network of both cohorts for CD; B) Abundance of the Bern clinics cohort overlaid with an integrated network of both cohorts for CD. C) The abundance of the Swiss IBD cohort overlaid with an integrated network of both cohorts for UC; D) Abundance of the Bern clinics cohort overlaid with an integrated network of both cohorts for UC.

### Comparative analysis

Figs. 3-5 illustrate the functionalities in MIND Web that can be used to readily compare distinct networks obtained from the same study. A comparison of networks across different studies, however, is more challenging for several reasons, including the fact that nodes in the corresponding networks may represent different taxonomy ranks. To address this problem, we developed a function named “level-up” that projects nodes (and corresponding edges) at a given level to a desired higher taxonomic rank (see details in Fig. S1). This is achieved by integrating lineage information from the NCBI taxonomy database as detailed in Fig. S6. During the level-up process, nodes belonging to a given taxonomic level (lower than the desired taxonomic rank) are merged into one node, and nodes with higher taxonomic ranking are unchanged. For example, two species nodes belonging to the same genus will be merged into one geneus node upon selecting the level-up function from species to genus; in this case, edges between the two species are ignored in the level-up process, while edges connecting the two species to other species are translated into edges between genus nodes (see additional explanations in Fig. S6). This implementation of the ”level-up” function is very simplistic, and could be substituted in the future by more sophisticated taxonomic projection estimations.

Fig. 6 shows an example of the application of the level-up function. In this example, we compared two independent studies that investigate bacterial vaginosis. The results of study 1 [68] involves 43 nodes with 900 edges (Fig. 6A), and the results of study 2 [69] involves 16 nodes with 120 edges (Fig. 6B). The merged network of both studies (Fig. 6C) indicates that the results of the two studies are inconsistent because only 23% of nodes and 6% of edges of study 1, and 69% nodes and 34% edges of study 2 overlap. However, visual examination of the network overlap (Fig. S7) suggest that correlations for overlapping edges agree relatively well: in particular the sign of correlation (positive or negative) in one study is highly conserved in the other. Using the level-up functions, we noticed that, at the Genus level, there are only two nodes specific to study 2 (labeled nodes in Fig. 6D); at Order level there is only one node specific to study 2 and three nodes specific to study 1 (Fig. 6E), and at the Phylum level study 2 turns into a subset of study 1 (Fig. 6F), suggesting that upon performing this coarse graining, the two studies display an overlapping interaction network topology.

**Figure 6.**
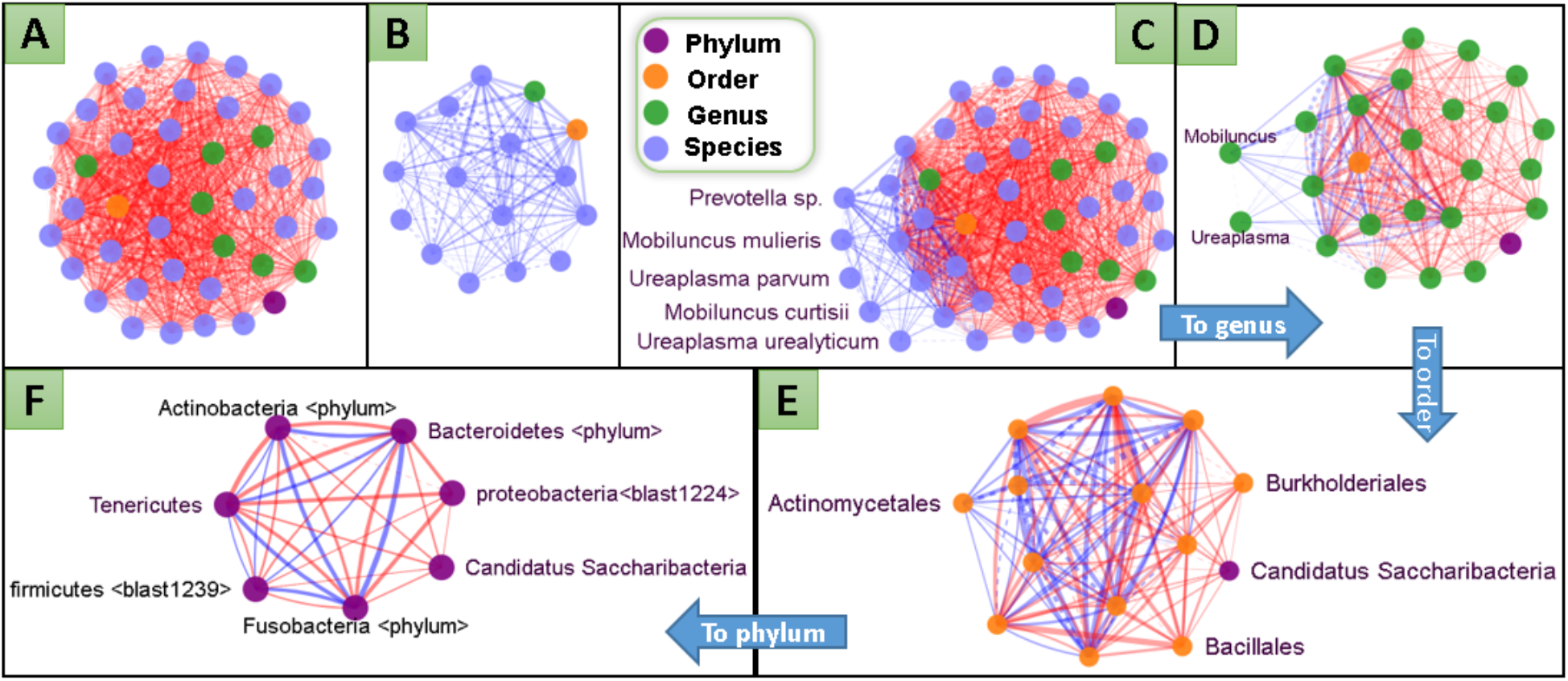
Comparison of the microbial network of bacterial vaginosis (BV) from two independent studies. The color of the nodes represents the taxonomic ranks, while the color of the edges represent different contexts. A) Study 1: vaginal microbial co-occurrence network created using Pearson correlation based on the 16 sRNA sequence data of 220 women (MIND context ID:C2000) [68]; B) Study 2: vaginal microbial co-occurrence network created using Spearman correlation based on the qPCR and PCR data of selected bacteria for 177 women (MIND context ID:C2002) [69]; C) Integration of two networks indicated that five microbes (left of the main cluster) are present only in Study 2; D) Upon using the “level-up” function to project all data to the Genus level, only two genera (Ureaplasma and Mobiluncus, all edges connected to them are in blue color only) specific to Study 2 were left; E) When we level-up to Order, there is only one node (Actinomycetales) that is specific to study 2, and three nodes specific to study 1; F) Upon leveling-up to Phylum, study 2 becomes a subset of study 1 with two phyla specific to study 1 (right side of the panel; these can be recognized because all edges that connect to them are orange).

### A case study for MIND: microbial networks associated with colorectal cancer

In order to illustrate the potential usage of MIND to perform complex multi-step comparisons, we reanalyzed a collection of datasets on microbial communities found in the gut mucosa and associated with colorectal carcinogenesis [70]. We used MIND to compare distinct datasets, and infer and interpret network modules. We applied very different approaches relative to the original publication [70], taking advantage of the functionalities provided by the MIND platform. A detailed description of the steps required to achieve the network shown in Fig. 7 can be found in the Supplementary Material. As shown in Fig. 7A-C, we first noticed that the network of normal mucosa has more nodes (43) and edges (132) than both the adenoma mucosal (38 nodes, 62 edges) and carcinoma mucosal (37 nodes and 78 edges) networks. This observation suggests that the normal mucosal microbial network is more diverse and more tightly coupled than microbial networks in transformed mucosal tissue.

**Figure 7.**
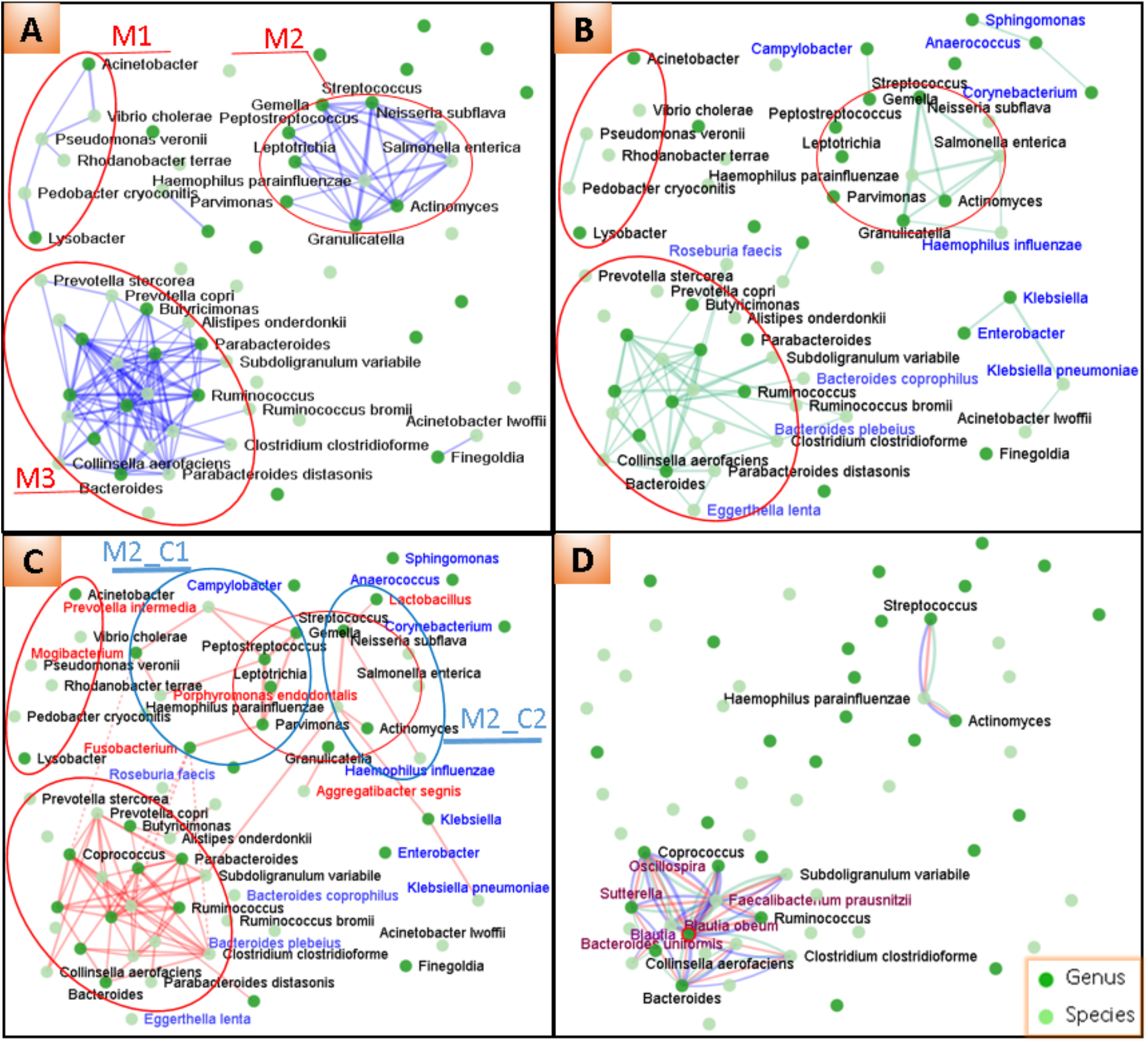
The modular structure of the correlation network of the gut mucosal microbiome across stages of colorectal carcinogenesis, based on data from [70]. A) The microbial correlation network of the normal gut mucosa in colorectal cancer (CRC) samples. Here, only nodes involved in modules M1, M2 and some of the nodes in M3 are labeled. In the microbial network of the adenoma (B) and carcinoma (C) mucosa in CRC samples, new nodes appear in the different modules, labeled respectively in blue and red. D) Overlapping networks of the normal, adenoma and carcinoma mucosa. Only connections that appear in all three contexts are shown and all nodes in the modules that are not labeled in Fig. 7A, 7B and 7C are labeled in purple.

We further noticed that the network of normal mucosa exhibits a clear modular structure, with three main modules (M1-M3) shown in Fig. 7A. Modules M1, M2 and M3 involve 6, 10 and 23 nodes respectively and they change significantly in the mucosa of adenoma and carcinoma to modules with a reduced number of members and connections (Fig. 7B-C). M1 has only two members left in the mucosa of adenoma, and disappears completely in the mucosa of carcinoma; M2 retains its core modular structure with five members left and one additional member (*Haemophilus influenzae*) in the adenoma mucosa. In the carcinoma mucosa, M2 is separated into two modules with the addition of some important members including *Fusobacterium* and *Prevotella intermedia* (Fig. 7C). The former has the species *Fusobacterium nucleatum* that were shown to be associated with CRC [71–73] and the latter has been found to be one of the leading causes of periodontal disease [74]; M3 is the most stable module with the majority of its members and connections remaining unchanged among three contexts (Fig. 7D).

Interestingly, modules M2 and M3 in the carcinoma are connected by *Haemophilus parainfluenzae* and *Subdoligranulum variabile* while M2 being spliced into two modules. The association between two microorganisms and CRC has been well reported [75–77]; its detailed mechanism however is not yet clear. This observation supports the hypothesis that the disruption of M2, and the interdependence between oral microbiota and gut ones, may play a role in the dysbiosis associated with CRC.

FInally, inspired by the recent report that the oral microbiome may provide clues to CRC [78], we decided to look into the potential connections between oral microbiome and networks in the CRC studies. We thus overlaid the CRC datasets with two networks summarizing known inter-bacterial interactions in the oral biofilm [79] (C0007-C0008 for early and later stage colonization). We found only two overlapping nodes (*Prevotella intermedia* and *Haemophilus parainfluenzae*), probably due to the fact that all nodes in the oral microbiome networks have species ranking while more than 50% of the nodes in the CRC studies have a genus ranking or higher. Interestingly, both nodes are part of the M2 module (either the original module for normal colon (Fig. 7A) or the extended M2 for the carcinoma (Fig. 7C)), pointing to possible hints for follow up studies.

## Discussion

Microbial interdependencies shape the structure, functions and dynamics of microbial communities. Physical and chemical interactions, mediated by a plethora of mechanisms, can modulate competition and cooperation, and hence the time and space-dependent abundance of microbes. In turn, from multiple types of data, one can infer correlations between organisms, which provide a pairwise readout of how the presence of one species can be the indicator of the presence or absence of another. Collectively, for a given type of microbial ecosystem (e.g., the gut microbiome under a given disease, or the rhizosphere community for a given plant), multiple heterogeneous datasets on interactions and correlations might be available, which convey complementary information about that community. MIND enables corroboration across datasets, and direct comparisons between correlations and interactions, with the potential to spark new understanding of how communities assemble and operate.

Some MIND functionalities, such as the comparative visualization of heterogeneous networks with different meanings and cutoffs, and the coarse graining based on the level-up function for taxonomy, should be viewed as a prototypes that will likely need further research and improvements in future iterations of microbial interaction databases and analysis tools. At the same time, the availability of such prototypes, and of our JSON format, could serve as a starting point for future conversations on how to best encode and integrate microbial interaction networks in a general and scalable way. Moreover, while the current version of MIND supports only pairwise interactions, there is no reason that the current formatting and representation could not be extended to higher order interactions.

Future releases of the MIND database and web platform could accommodate a broader range of microbial networks to cover more soil and marine microbiome, as well as disease and animal-associated microbiome datasets. When integrated with modeling and simulation tools such as COMETS [80], our platform will also be able to perform analyses of natural and artificial microbial communities, and could thereby become a powerful tool for personalized biomedicine, metabolic engineering, and biogeochemistry. In the future, by extending to MIND the metagraph technology that was previously developed for VisANT, it would be possible to encode both intracellular microbial circuits, and inter-species interactions, effectively enabling multi-scale representation of knowledge on microbial metabolism and interactions from individual enzymes to ecosystems.

## MATERIALS AND METHODS

### Design and implementation of MIND Database

To encode interaction networks under a given set of conditions, we adopted a context-based data structure, as shown in Fig. S5. Each interaction network is organized by a specific Context that stores abundant metadata, including information about the environment, disease relevance, and experimental and computational methods used to infer interactions.

The MIND database uses the NCBI taxonomy ID [58,59] to represent each microbe involved in the interactions. As different datasets incorporated in the database typically report interactions at different taxonomic levels, it is also crucial to include the taxonomy hierarchy in the database. From this perspective, NCBI taxonomy is integrated into MIND and periodically synchronized to keep the data up to date. Table S2 provides a brief summary of the data currently available in the MIND database.

### System Architecture

To ensure scalability, extensibility, and interoperability, the MIND system adopts a service-oriented architecture (SOA) [53] that is based on VisANT-Predictome’s three-tier system [55,57]. The architecture is based on a collection of web services each of which carries out functions that are well-defined, self-contained, and independent of the context or state of other services, with client-service protocols translated via an enterprise service bus (ESB). This facilitates the integration of different architectural systems, regardless of whether the services are internal or external. In combination with the fact that the web-based application itself is independent of the operating system, such architecture achieves substantial interoperability and facilitates easier integration of different architectural systems regardless of whether the services are internal or external, therefore allowing us to draw upon analytical methods developed by ourselves and others.

PostgreSQL is used to store all data related to interactions and networks. The web platform is built using the latest web technology and implemented as a Single Page Application (SPA). It, therefore, can be accessed wherever the browser is available while providing rich functionalities similar to the Desktop application. The web platform uses a list of open source libraries to speed up the development, with a Bootstrap framework to make the platform mobile friendly, AngularJS [64] to improve the user interactivity and D3 [62] to enhance the visualization. All web services, as well as the ESB, are implemented using J2EE technology and hosted by Apache Tomcat.

### Data Sources

The data currently available as default in the MIND database come from two main sources: i) curated manually from the networks in published studies, ii) generated by the MiCONE pipeline (see [23]) with recommended settings based on published 16s rRNA sequence data.

#### Data curation

It is difficult in general to integrate or compare microbial networks published in the literature because the nodes of these networks are often represented by OTUs that are not comparable across different studies. The curation of these networks however not only enables integrative or comparative studies but also enhances our understanding of the mechanisms by which microbial cells interact with one another and with host cells. In addition to correlation networks, our set of curated data includes networks of interactions resulting from evolutionary studies such as lateral gene transfer, metabolic influence, as well as modeling and simulations by tools such as COMETS [80]. Other than the conversion of OTUs to the NCBI taxonomy IDs, the networks we curated are kept unchanged from the original publications: the additional modification or integration can be performed at the client side as per users’ choices. In this way, the platform provides great flexibility while retaining the data originality.

### Standardization and Data Provenance

During the development of MIND 1.0, we carefully evaluated the current standards for interactions and networks, including the HUPO PSI’s molecular interaction format (PSI-MI) [81], Biological Pathways Exchange format (BioPAX) [82], and Systems Biology Markup Language (SBML) [83], as well as ones for the metadata in the metagenomic studies, such as the minimum information about a genome sequence (MIGS) [84]. We then developed a simplified JSON format that combines both the network and metadata information to facilitate the submission of user-generated networks. We use context to represent all the metadata associated with the network and each network in MIND database is assigned a unique context ID. The format may evolve based on feedback from the community in the future. An important part of the JSON format [85] is the metadata specified for the data provenance, including the PubMed ID and data of the publication, as well as the email of corresponding authors or submitters so that users can directly communicate with them when questions arise. All networks stored in the MIND database can be exported in MIND data JSON formats; and any networks in MIND data JSON format can be loaded in MIND-web through drag & drop operations.

### Functionalities of the MIND Web Platform

#### Import and Output

Networks shown in the platform can be imported/exported as MIND JSON data format (http://www.microbialnet.org/mind/mindweb/formats_full.html) as mentioned earlier, as well as MIND extended JSON format which saves the network information together with all the current configurations of the platform (such as edge weight cutoff, etc., similar to VisML). The network may also be exported as a PNG image and SVG graph (Fig. S1).

#### User Registration

The MIND platform is free to use without user registration. Registration however is encouraged because it not only allows user to receive the latest update of the database but also to use a list of functions that only available after user’s login, including online saving/loading of networks to/from the server, sharing networks to the colleagues, submitting the network to the MIND database directly. Registered VisANT users are automatically enrolled in MIND platform as MIND and VisANT share an overlapping user base.

## Acknowledgments

This work was supported by the National Institutes of Health, R01GM121950 to DS, ZH and CD. DS further acknowledges support by the NSF Center for Chemical Currencies of a Microbial Planet (CCoMP), and the NIH National Institute on Aging, award number UH2AG064704.

## Authors’ contributions

DS, ZH and CD directed the project. ZH and DS conceived the platform structure. ZH designed the software architecture. ZH and YW implement the platform. ZH and DK collected the data and performed the analysis. GB tested the platform and contributed to the computational analysis. KK contributed to the overall development of the project. ZH and DS prepared the manuscript and all authors contributed the writing.

