## Supplementary Material for "A resource for the comparison and integration of heterogeneous microbiome networks"

### Supplementary figures

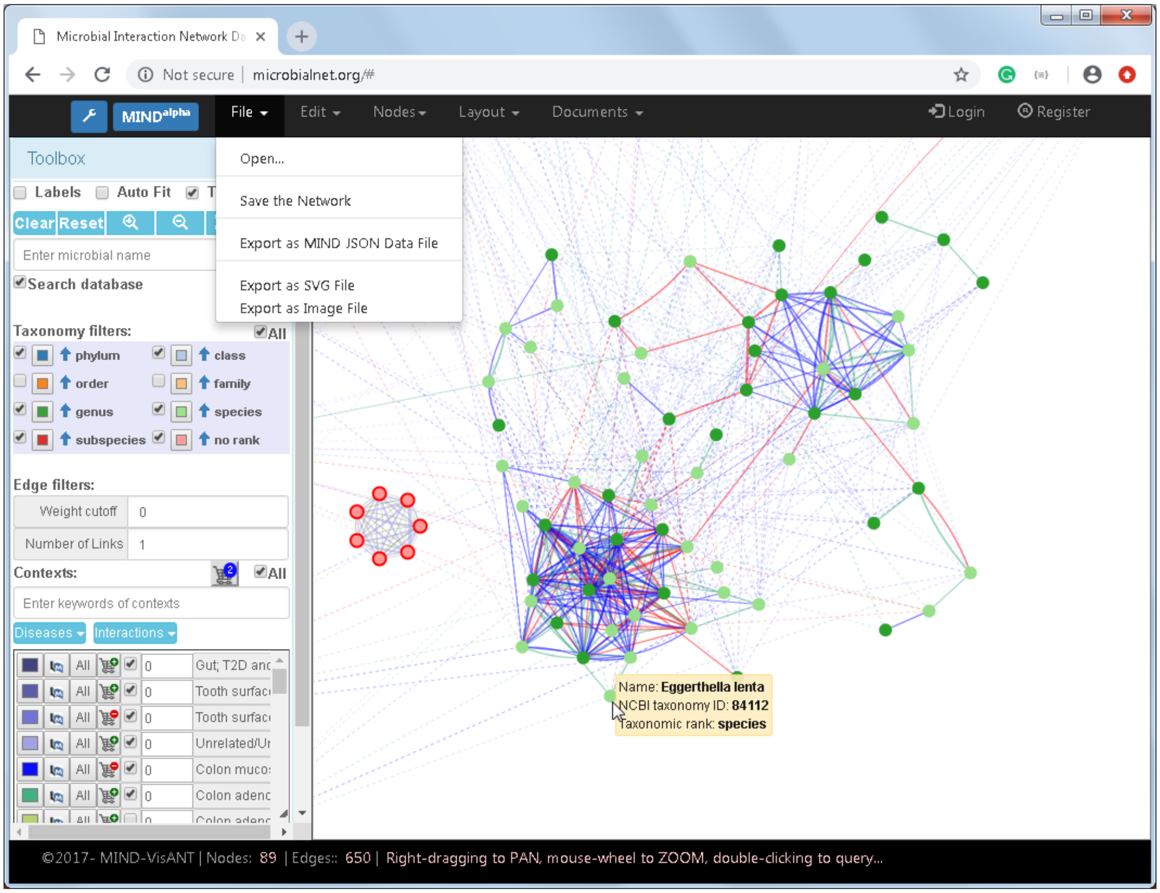

**Fig. S1.** Overview of the MIND Web platform. The top is the menu bar; the bottom is the status bar. The left is the toolbox, and the rest of the space is the network panel.

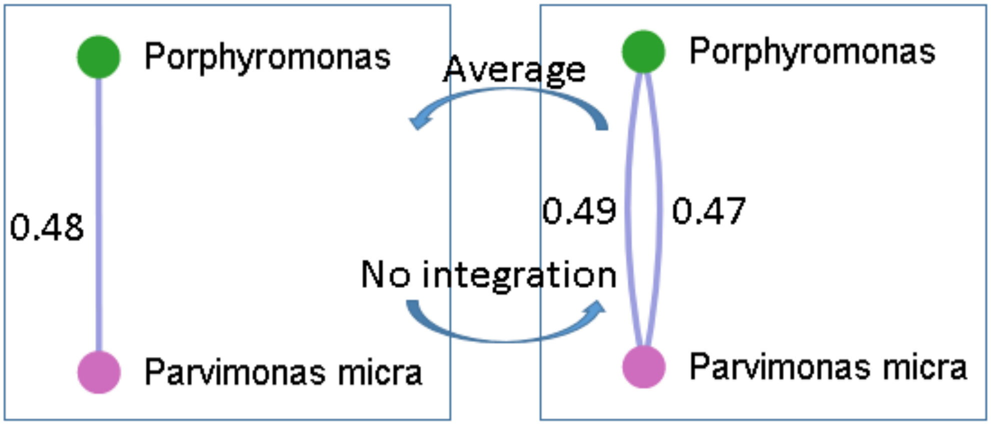

**Fig. S2.** Multiple interactions may be present in the same context for a given pair of microbes.

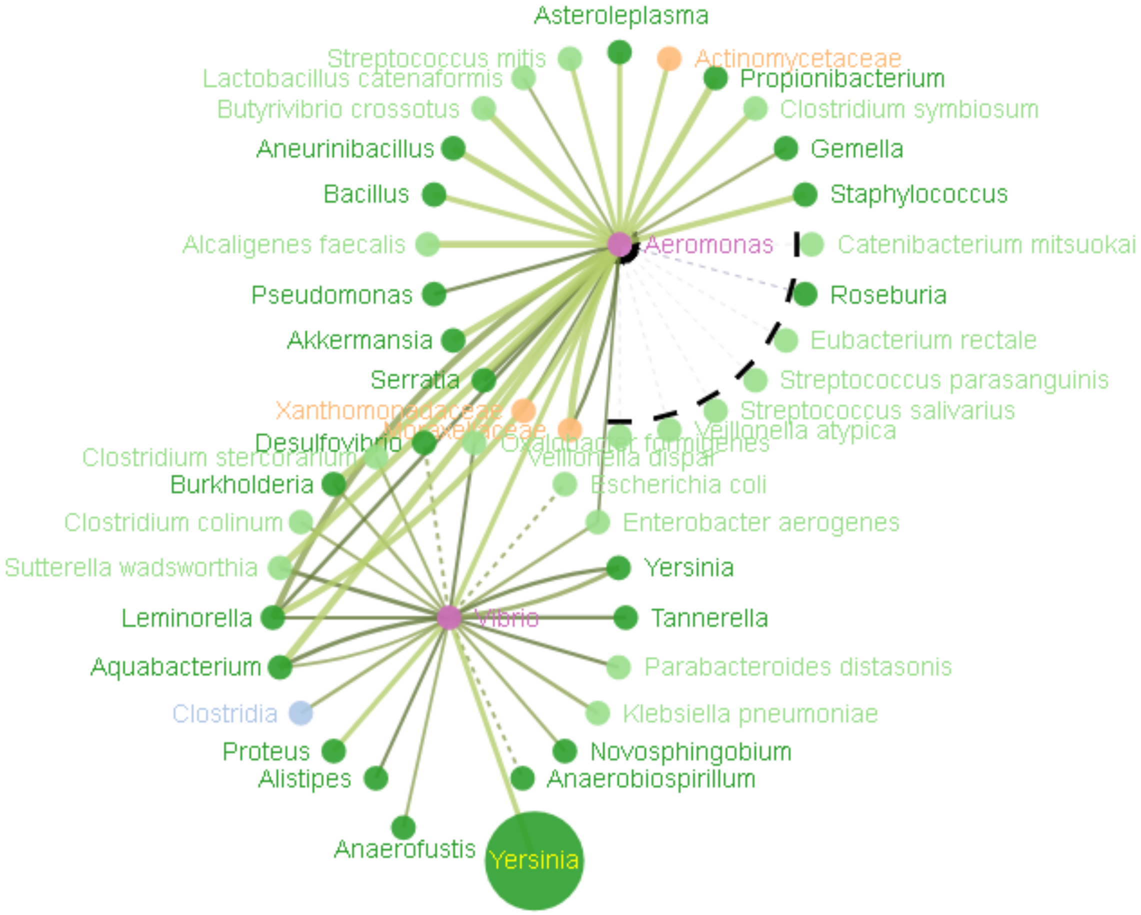

**Fig. S3**. Illustration of the optimized label arrangement and customization. Except for the nodes associated with *Anaerofustis* and *Yersinia*, the labels of the other nodes are positioned automatically.

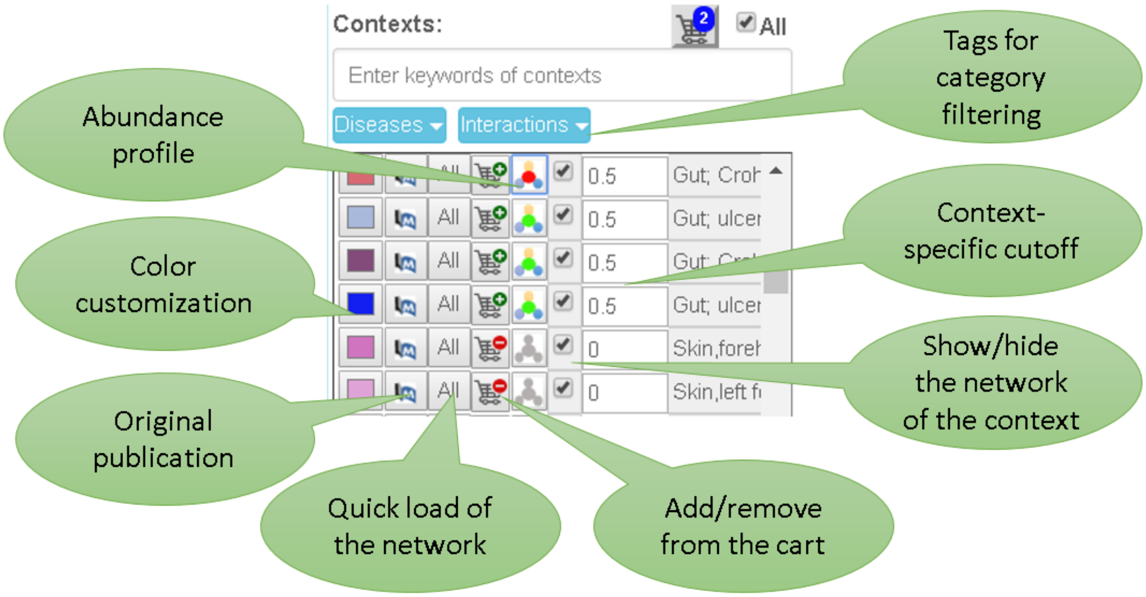

**Fig. S4**. Details of the MIND dataset navigation panel, where functions associated with context navigation are displayed. In addition to filtering contexts by different criteria (light blue buttons on top), one can, for each dataset: (1) Select the edge color; (2) Visualize the publication associated with the data; (3) Load the network; (4) Add/Remove the dataset rom the cart (to enable selection and handling of multiple datasets); (5) Switch on/off taxonomic abundane profile; (6) Show/hide the network; (7) Select a network specific edge weight cutoff; and (8) View keyworkds associated with the context.

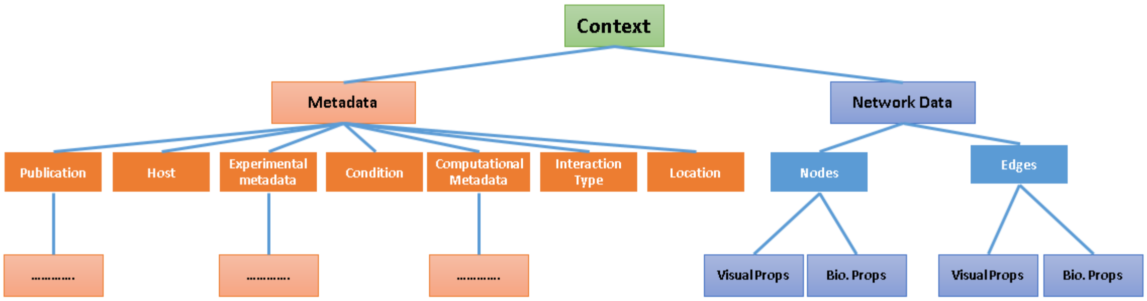

**Fig. S5**. Data organization in the MIND platform. Each network is represented by a given context where a set of flexibly structured metadata describe the conditions under which the network is curated. Visual properties of both nodes and edges are distinct from the biological properties required by the Model-View-Control design pattern. This architecture facilitates the support of metagraphs [18], which in the future could be used to embed module information (such as taxonomy/disease hierarchy) into the network.

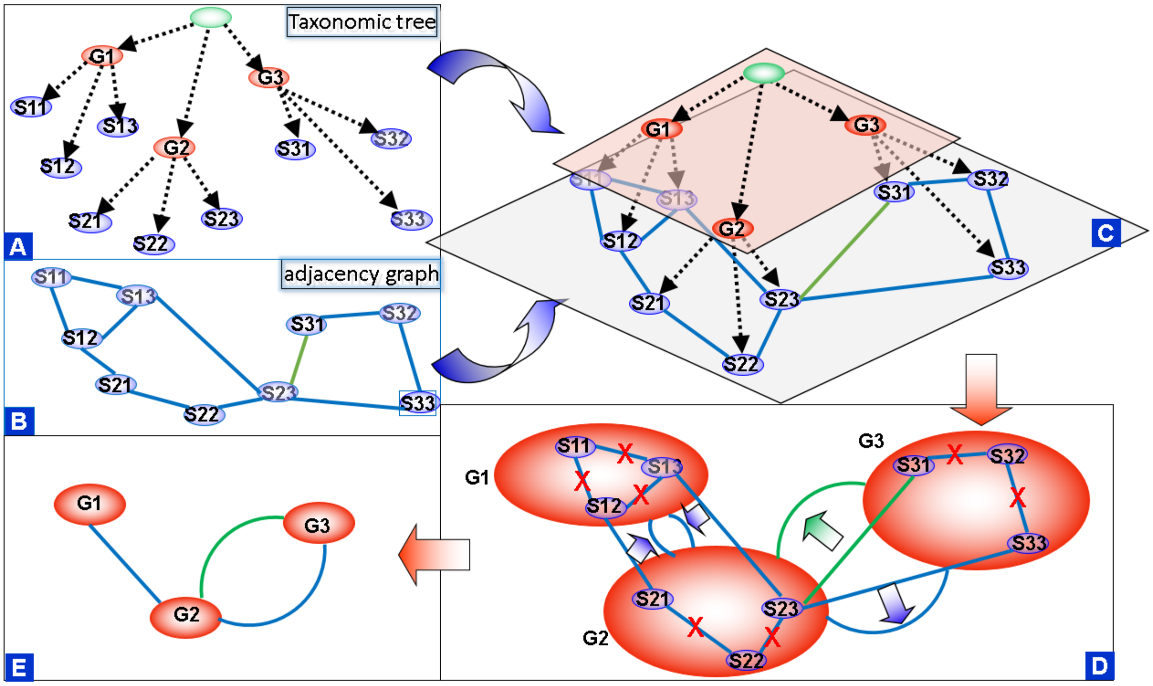

**Fig. S6**. Illustration of taxonomic level-up approximation from species to genus. A) Partial axonomy tree where there three geneus nodes each have three species nodes. B) A network between nine species nodes where different edge color represents different contexts. C) Integration of the taxonomy tree and the network in 3D space. D) In projecting the 3D integrated graph into 2D space, all inter-genus links are ignored, and intra-genus links are transferred to the genus nodes. E) The network between genus nodes after level-up approximation; by default, links of the same context between a given node pair are integrated with their weight averaged (Fig. S2) (a step which is reversible).

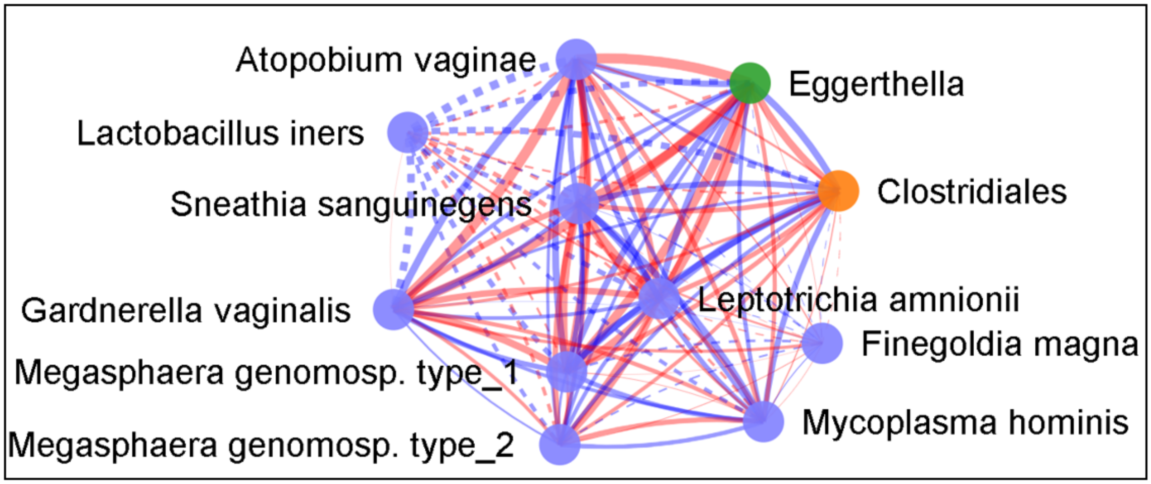

**Fig. S7.** Overlapping of the networks between two independent studies of bacterial vaginosis.

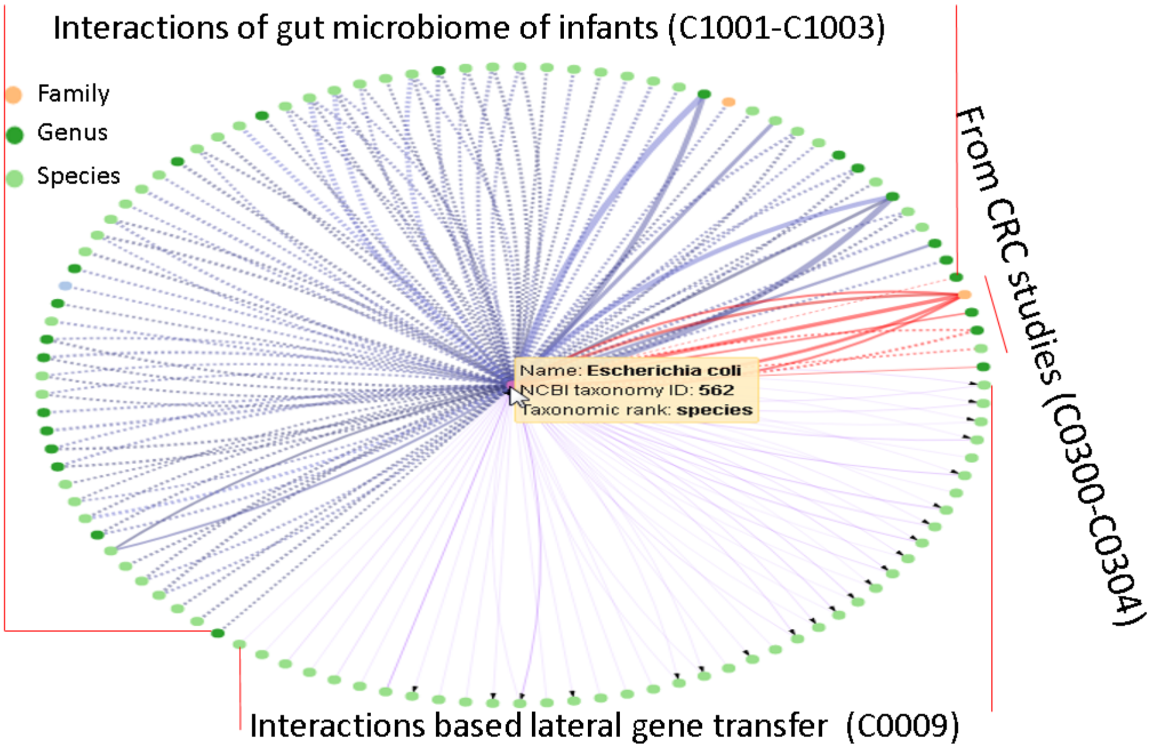

**Fig. S8.** Microorganisms that interact with *E. coli* are queried against the MIND database. The blue lines indicate the correlation edges (Pearson correlation, stool samples) of gut microbes in new-born babies delivered either vaginally or by at-term cesarean (context ID C1001-C1003)[4]. The red lines represent the co-occurrence correlation of gut mucosal microbiome identified by SparCC [19] in a study of colorectal cancer (C0300-C0304) [5]. The purple lines correspond to genes that were transferred between bacterial genomes via a phage where the arrow indicates the direction of the transfer events [6]. The dashed line represents the negative correlation while the solid line represents the positive one.

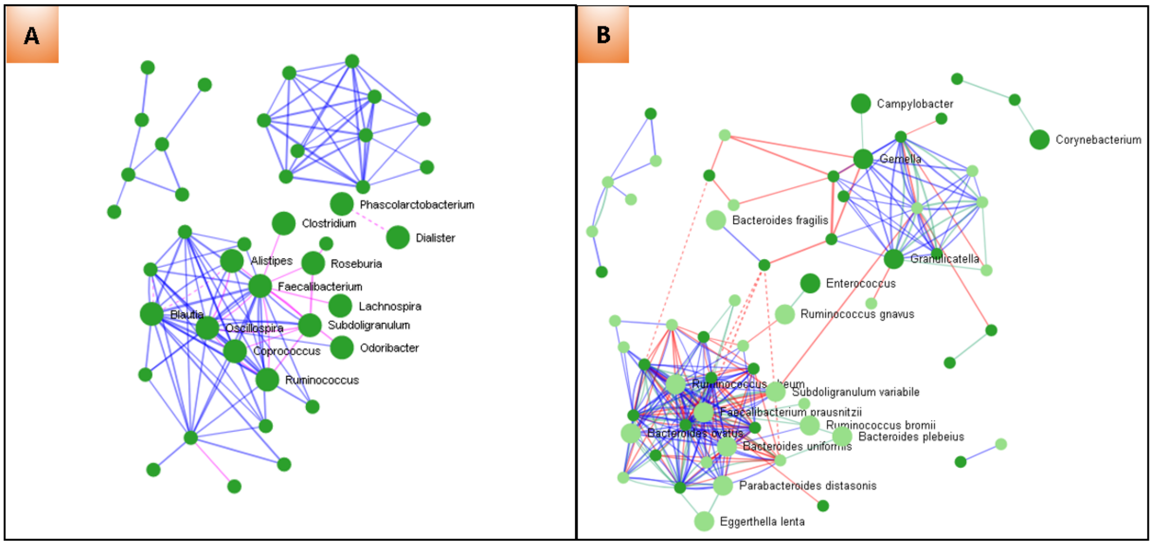

**Fig. S9.** Comparison of the network of gut microbiome of healthy samples against CRC studies. The overlapping nodes are labeled and shown with bigger size. While CRC studies use the mucosa samples, the microbiome of the healthy gut are all based on stool samples. A) Comparison of the microbial network of the normal gut mucosa in CRC samples (blue lines) against of the network of gut microbial network of the healthy samples (stool) (purple lines) with original data reported in the HMP 1 project [20] and network data reproduced using MIND pipeline. B) Comparison of networks of gut microbiome of new born babies (1-3 days after the deliver, either at-term caesarean or vaginally) [4].

### Supplementary tables

| Table S1 Constitution of module M2 | | | |
|---|---|---|---|
| Name | Tax ID, rank | Habitat | Disease-associated |
| *Peptostreptococcus* | 1257, genus | mouth, skin, gastrointestinal, vagina and urinary tracts | endocarditis[21,22], paravalvular abscess, pericarditis, CRC[8–10] |
| *Haemophilus parainfluenzae* | 729, species | part of the normal flora of the mouth | Endocarditis[23,24], dental abscesse |
| Streptococcus | 1301, genus | mouth | endocarditis, pneumonia, meningitis |
| Gemella | 1378, genus | mouth | Endocarditis, CRC[8–10] |
| Actinomyces | 1654, genus | Mouth, skin, gut | Periodontitis, endocarditis |
| Neisseria subflava | 28449, species | Mouth, upper respiratory tract | meningitis, septicemia endocarditis [25] |
| Salmonella enterica | 28901, species | Not human microbiota?? | among the main causes of bacterial gastrointestinal infections. The highest infection risk is oral ingestion of contaminated food or water[26] |
| Leptotrichia | 32067, genus | Mouth, gut | periodontal diseases and oral cavity abscesses, endocarditis[11][12], CRC[27,28] |
| Granulicatella | 117563, genus | Mouth | dental plaque , dental abscesses, endodontic infection[29], endocarditis[30][29] |
| Parvimonas | 543311, genus | Mouth | endocarditis [31], CRC[8–10] |
| Haemophilus influenzae | 727, species | Not human microbiota?? | endocarditis (bacteremia)[17] |
| Campylobacter | 194, genus | Mouth | periodontal diseases and oral cavity abscesses, endocarditis[14–16], CRC[27,28] |
| Fusobacterium | 848, genus | Mouth, gut | Periodontitis, endocarditis [32], CRC[8–10] |
| Prevotella intermedia | 2831, species | Mouth | Gingivitis, Periodontitis, |
| Mogibacterium | 86331, genus | Mouth, | Periodontitis[33] |
| Porphyromonas endodontalis | 28124, species | Mouth | Periodontitis[34] |
| Lactobacillus | 1578, genus | Mouth | dental caries, periodontitis[35] endocarditis[36,37] |
| Aggregatibacter segnis | 739, species | Mouth | endocarditis[38] |

| | | | |
|---|---|---|---|
| Klebsiella pneumoniae | 573, species | Gut | endocarditis[39,40] |
| Klebsiella | 570, genus | Gut | endocarditis [41,42] |
| Haemophilus influenzae | 727, species | Not human microbiota | endocarditis (bacteremia)[17] |

**Table S2: List of contexts available in MIND database** [48 contexts involving 448847 interactions between 4142 microbes] (by 4/10/2019)

| Context ID | Location | Interaction type | Disease state | Condition | PubMed ID |
|---|---|---|---|---|---|
| C0008 | Tooth surface | binding | Healthy | Late colonizers | 12209001 |
| C0007 | Tooth surface | binding | Healthy | Early colonizers | 12209001 |
| C3003 | Attached Keratinized gingiva | correlation | Healthy | 242 healthy adults (129 males, 113 females) | 22699609 |
| C3004 | Buccal mucosa | correlation | Healthy | 242 healthy adults (129 males, 113 females) | 22699609 |
| C3006 | Left antecubital fossa | correlation | Healthy | 242 healthy adults (129 males, 113 females) | 22699609 |
| C0500 | Skin forehead | correlation | Healthy | 200 skin samples from Chinese individuals living in Hong Kong | 26177982 |
| C1000 | Adenoid | correlation | Unrelated/ Uncertain | 67 individuals who underwent adenoidectomy, weight=MIC-??2 > 0.2 | 23113966 |
| C3001 | Mid vagina | correlation | Healthy | 242 healthy adults (129 males, 113 females) | 22699609 |
| C0302 | Colon adenoma-adjacent mucosae | correlation | Colon adenoma | Gut mucosal microbiome of 47 paired samples of adenoma and adenoma-adjacent mucosae | 26515465 |
| C0303 | Colon carcinoma mucosa | correlation | Colon carcinoma | Gut mucosal microbiome of 52 paired samples of carcinoma and carcinoma-adjacent mucosae | 26515465 |
| C0304 | Colon carcinoma-adjacent mucosa | correlation | Colon carcinoma | Gut mucosal microbiome of 52 paired samples of carcinoma and carcinoma-adjacent mucosae | 26515465 |
| C0300 | Colon mucosae | correlation | Normal | Gut mucosal microbiome of 61 healthy controls | 26515465 |
| C0301 | Colon adenoma mucosae | correlation | Colon adenoma | Gut mucosal microbiome of 47 paired samples of adenoma and adenoma-adjacent mucosae | 26515465 |
| C0305 | Skin axilla | correlation | illary odour | 24 Caucasian male and female non-antiperspirant volunteers | 25653852 |
| C0306 | Gut | correlation | Healthy | 41 healthy controls, 56 subjects with prehypertension, and 99 patients with primary hypertension, northern China, newly diagnosed hypertensive patients prior to antihypertensive treatment,stool sample | 28143587 |
| C0307 | Gut | correlation | Healthy | 41 healthy controls, 56 subjects with prehypertension, and 99 patients with primary hypertension, northern China, newly diagnosed hypertensive patients prior to antihypertensive treatment,stool sample. | 28143587 |
| C2000 | Vagina | correlation | Bacterial vaginosis | 220 women, 34% were Black and 44% were White women, 98 (43%) had BV by Amsel?€?s criteria, and 117 (53%) by Gram stain | 22719852 |

| | | | | | |
|---|---|---|---|---|---|
| C0308 | Gut | correlation | Prehypertension | 41 healthy controls, 56 subjects with prehypertension, and 99 patients with primary hypertension, northern China, newly diagnosed hypertensive patients prior to antihypertensive treatment,stool sample. | 28143587 |
| C0309 | Gut | correlation | Hypertension | 41 healthy controls, 56 subjects with prehypertension, and 99 patients with primary hypertension, northern China, newly diagnosed hypertensive patients prior to antihypertensive treatment,stool sample. | 28143587 |
| C2001 | Gut | correlation | Healthy | 96 healthy individuals who were previously characterized for their bacteria/diet relationships | 23799070 |
| C2002 | Vagina | correlation | Bacterial vaginosis | Self-collected vaginal smears and swabs were obtained from 177 women | 24131550 |
| C3007 | Left Retroauricular crease | correlation | Healthy | 242 healthy adults (129 males, 113 females) | 22699609 |
| C3008 | Palatine Tonsils | correlation | Healthy | 242 healthy adults (129 males, 113 females) | 22699609 |
| C3010 | Right Antecubital fossa | correlation | Healthy | 242 healthy adults (129 males, 113 females) | 22699609 |
| C3011 | Right Retroauricular crease | correlation | Healthy | 242 healthy adults (129 males, 113 females) | 22699609 |
| C3013 | Stool | correlation | Healthy | 242 healthy adults (129 males, 113 females) | 22699609 |
| C3014 | Subgingival plaque | correlation | Healthy | 242 healthy adults (129 males, 113 females) | 22699609 |
| C3016 | Throat | correlation | Healthy | 242 healthy adults (129 males, 113 females) | 22699609 |
| C3017 | Tongue dorsum | correlation | Healthy | 242 healthy adults (129 males, 113 females) | 22699609 |
| C3005 | Hard palate | correlation | Healthy | 242 healthy adults (129 males, 113 females) | 22699609 |
| C3009 | Posterior fornix | correlation | Healthy | 242 healthy adults (129 males, 113 females) | 22699609 |
| C3012 | Saliva | correlation | Healthy | 242 healthy adults (129 males, 113 females) | 22699609 |
| C3015 | Supragingival plaque | correlation | Healthy | 242 healthy adults (129 males, 113 females) | 22699609 |
| C3002 | Anterior nares | correlation | Healthy | 242 healthy adults (129 males, 113 females) | 22699609 |
| C3018 | Vaginal introitus | correlation | Healthy | 242 healthy adults (129 males, 113 females) | 22699609 |
| C0502 | Skin right forearm | correlation | Healthy | 200 skin samples from Chinese individuals living in Hong Kong | 26177982 |
| C0503 | Skin left palm | correlation | Healthy | 200 skin samples from Chinese individuals living in Hong Kong | 26177982 |
| C0504 | Skin right palm | correlation | Healthy | 200 skin samples from Chinese individuals living in Hong Kong | 26177982 |
| C0501 | Skin left forearm | correlation | Healthy | 200 skin samples from Chinese individuals living in Hong Kong | 26177982 |
| C1001 | Gut | correlation | Healthy | CS1-3 following birth, total 25 at-term caesarean (CS) delivered neonates, stool sample | 26332837 |
| C1002 | Gut | correlation | Healthy | CS7-30 following birth, total 25 at-term caesarean (CS)delivered neonates, stool sample | 26332837 |
| C1003 | Gut | correlation | Healthy | V1-3 days following birth, total 6 vaginally (V) delivered neonates, stool sample | 26332837 |
| C1802 | Gut | metabolic competition index | Mixed | the composition of naturally occurring communities as measured by metagenomic sequencing of faecal specimens from 124 healthy, overweight and obese individual human adults, as well as | 23858463 |

| | | | | | |
|---|---|---|---|---|---|
| | | | | inflammatory bowel disease (IBD) patients, from Denmark and Spain | |
| C1803 | Gut | metabolic competition index | Mixed | the composition of naturally occurring communities as measured by metagenomic sequencing of faecal specimens from 124 healthy, overweight and obese individual human adults, as well as inflammatory bowel disease (IBD) patients, from Denmark and Spain | 23858463 |
| C1800 | Tooth surface | metabolic complementarity index | Healthy | oral species known to influence one another?€?s growth in shared environment, ranging from initial colonizers to late-arriving pathogens | 23858463 |
| C1801 | Tooth surface | metabolic complementarity index | Healthy | oral species known to influence one another?€?s growth in shared environment, ranging from initial colonizers to late-arriving pathogens | 23858463 |
| C0006 | Gut | metabolic influence | T2D and non-diabetic control | male, mid-age and normal-weight cohort | 28585563 |
| C0009 | | transduction | Unrelated/ Uncertain | lateral gene transfer by transduction from donors → recipients mediated by the phages | 27648812 |

### Supplementary Text

#### Steps to achieve the network shown in Fig. 5

Figure 5A-C display the weightad network with cutoff 0.25 for normal mucosa (MIND context ID C0300), adenoma mucosa (C0301) and carcinoma mucosa (C0303) respectively. The 0.25 cutoff is chosen because all negative correlations (shown as dashed lines) in the normal mucosa disappear with this threshold (the original publication used 0.3 as the cutoff [5]).

#### Features of module 2 and its change across the different stage of CRC

<u>Structure of module M2 in the mucosa of the normal colon</u>. Except for *Salmonella enterica*, all taxa are part of oral microbiota according to the Human Oral Microbiome Database (HOMD) [7], and many of them are associated with gum diseases such as periodontitis.  Others, such as *Peptostreptococcus* [8–10] and *Leptotrichia* [11,12], are being reported elsewhere to be associated with CRC. However, it is surprising that all the oral microbiota involved in M2 have been reported being involved in infective endocarditis: a disease that affects multiple systems and results from infection, usually bacterial, of the endocardial surface of the heart [13].

<u>M2 in the mucosa of adenoma</u>. Besides for members from M2 in the normal colon, there are new members: *Campylobacter* and *Haemophilus influenza* (Fig. 6B). The former is part of oral microbiota and are correlated with CRC[11,12], both however may lead to endocarditis [14–16][17]. Interestingly, *Salmonella enterica*, not a member of oral microbiota, is the member that has the most connections.

<u>M2 in the mucosa of carcinoma</u>. As shown in Fig. 6C, M2 in the carcinoma has been divided into two modules: M2_C1 and M2_C2. The former have four additional members: *Fusobacterium, Prevotella intermedia, Mogibacterium* and *Porphyromonas endodontalis* and the later have three new members: *Haemophilus influenza*, Lactobacillus, *Klebsiella pneumoniae* and Klebsiella. Among the new members, *Klebsiella pneumoniae* and *Klebsiella* are part of gut microbiota and the rest all are all part of oral microbiota except *Haemophilus influenzae*. All new members in M2_C1 are involved in periodontitis whereas fusobacterium has an important contribution to CRC. All new members of M2_C2, together with *Fusobacterium*, are associated with endocarditis.

33. Casarin RCV, Saito D, Santos VR, Pimentel SP, Duarte PM, Casati MZ, et al. Detection of Mogibacterium timidum in subgingival biofilm of aggressive and nondiabetic

and diabetic chronic periodontitis patients. Brazilian J. Microbiol. 2012;

34. Bedran TBL, Marcantonio RAC, Neto RS, Mayer MPA, Grenier D, Spolidorio LC, et al. Porphyromonas endodontalis in chronic periodontitis: A clinical and microbiological cross-sectional study. J. Oral Microbiol. 2012;

35. Kõll-Klais P, Mändar R, Leibur E, Marcotte H, Hammarström L, Mikelsaar M. Oral lactobacilli in chronic periodontitis and periodontal health: Species composition and antimicrobial activity. Oral Microbiol. Immunol. 2005;

36. Husni RN, Gordon SM, Washington JA, Longworth DL. Lactobacillus bacteremia and endocarditis: review of 45 cases. Clin. Infect. Dis. 1997;

37. Yagi S, Akaike M, Fujimura M, Ise T, Yoshida S, Sumitomo Y, et al. Infective Endocarditis Caused by Lactobacillus. Intern. Med. 2008;

38. Revest M, Egmann G, Cattoir V, Tattevin P. HACEK endocarditis: state-of-the-art. Expert Rev. Anti. Infect. Ther. 2016.

39. Pai RK, Wall TS, Macgregor JF, Abedin M, Freedman RA. Klebsiella pneumoniae: A rare cause of device-associated endocarditis. PACE - Pacing Clin. Electrophysiol. 2006;

40. Thomas MG, Rowland-Jones S, Smyth E. Klebsiella pneumoniae endocarditis. J. R. Soc. Med. 1989;

41. Anderson MJ, Janoff EN. Klebsiella Endocarditis: Report of Two Cases and Review. Clin. Infect. Dis. 1998;

42. Brouqui P, Raoult D. Endocarditis due to rare and fastidious bacteria. Clin. Microbiol. Rev. 2001.
